# An optimised computational approach for the identification of somatic structural variants in cancer

**DOI:** 10.1101/2025.07.01.662575

**Authors:** Sara Waise, Nana Mensah, Tom Lesluyes, Jonas Demeulemeester, Adrienne Flanagan, Nischalan Pillay, Peter Van Loo

## Abstract

Structural variants play a critical role in tumorigenesis. At present, these events are most commonly identified using short-read whole-genome sequencing data, and a number of computational tools are available for this purpose. Consensus approaches have been used to improve precision, but may reduce sensitivity. The optimal number and combination of callers remains unclear, in part due to the lack of gold standard real-world datasets for validation. Here, we benchmark the performance of Delly, GRIDSS, LUMPY, Manta and SvABA, using a validation set of consensus calls from the Pan-Cancer Analysis of Whole Genomes Consortium. Manta showed the best standalone performance, identifying 88% of the validation set calls, and was included in all of the best-performing caller combinations. A consensus approach comprising Delly, GRIDSS, Manta and SvABA was selected as the optimum approach from those tested. We provide a NextFlow implementation of our optimised consensus approach as a resource for the cancer genomics community.

## Introduction

Somatic structural variants (SVs) are a key type of mutation in human cancer^1,2^. These alterations may occur as simple events (such as insertions, deletions, duplications and translocations) or as part of more complex phenomena (e.g. chromothripsis, characterised by chromosome shattering with associated massive genomic rearrangements)^3-6^. To date, short-read whole-genome sequencing remains the most widely used genomic profiling method; emerging technologies such as optical mapping and long-read sequencing enable the detection of SVs human tissue more directly^7^. Due in large part to limitations of short-read whole genome sequencing, structural rearrangements remain difficult to characterise in such data^2,8,9^. However, a recent study suggests that an optimised somatic SV calling approach in short-read data can yield similar results to long-read sequencing when applied to regions of the genome mappable by short-read sequencing^10^.

A number of algorithms are available to call SVs in short-read WGS data^2^. These approaches rely on one or a combination of methods to identify rearrangements: split reads, discordant paired-end reads, read depth, and local assembly^9,11^. An ideal SV caller would have low false discovery rate and high accuracy^9^. In practice, each approach has relative strengths and weaknesses^2,5^, and no single caller consistently identifies all types and sizes of SVs^12^. Callers each employ different combinations of principles, and as a result there is rarely complete agreement between different algorithms^2^. Furthermore, breakpoints identified by SV callers may be considered imprecise, further complicating attempts to match variant calls between different algorithms^13^.

Increasingly, studies have used an ensemble approach for variant identification, with the aim of improving sensitivity and specificity^5,9,14,15^. This can improve performance relative to individual callers, but may, to some extent, sacrifice recall for higher precision^5^. The effectiveness of ensemble calling is highly dependent on the chosen callers: inappropriate selection can lead to reduced sensitivity or an increased rate of false-positive calls^2,5,9^.

Multiple tools for ensemble SV calling are now available, including MetaSV^16^, FusorSV^17^ and ConsensuSV^7^. However, the optimal number and combination of tools for consensus SV identification has not yet been determined. Previous studies advocate inclusion of assembly-based callers (*e*.*g*. Manta^18^, GRIDSS^19^) and those that incorporate multiple evidence sources^2,9^. Benchmarking of somatic SV calling is difficult due to the lack of gold-standard reference datasets^4,9^. Previous studies validating SV calling have predominantly relied on germline data, cell line data, or simulations, which do not capture the broader complexity of structural variants derived from tumour samples^2,4,5,9,16^.

Here, we present a benchmarking of somatic SV calling, using patient-derived data^20^ and five callers with documented precision^2^. We find that a consensus approach using variants identified by at least two out of four callers (Delly^21^, GRIDSS^19^, Manta^18^ and SvABA^22^) yields the best combination of sensitivity and minimised excess calls. The optimised approach, incoportating updated tools, somatic and panel of normal filtering and consensus calling, is made available as a Nextflow pipeline.

## Results

### Individual caller performance

To identify the number of somatic structural variants called by five specialised callers (Delly, v0.8.5, GRIDSS as gridss-purple-lix v1.3.2, LUMPY as Smoove v0.2.6, Manta, v1.5 and SvABA v1.1.0), we ran each tool individually with the developer-recommended settings on a set of 40 whole-genome sequenced tumour-normal pairs from the Pan-Cancer Analysis of Whole Genomes Consortium (PCAWG). These data were realigned to hg38 using the Genomics England Dragen 2.0 pipeline. We observed wide variation in the number of SVs identified by each caller (Fig. 1A). LUMPY made the highest median number of calls (538, IQR 410.5- 992.75), followed by Delly (median 493.5, IQR 275-1185) and GRIDSS (median 481, IQR 401- 784.25) SvABA and Manta made fewer calls overall, returning a median of 202 (IQR 95-798) and 77 (IQR 62.75-546.25) SVs, respectively.

**Figure 1.**
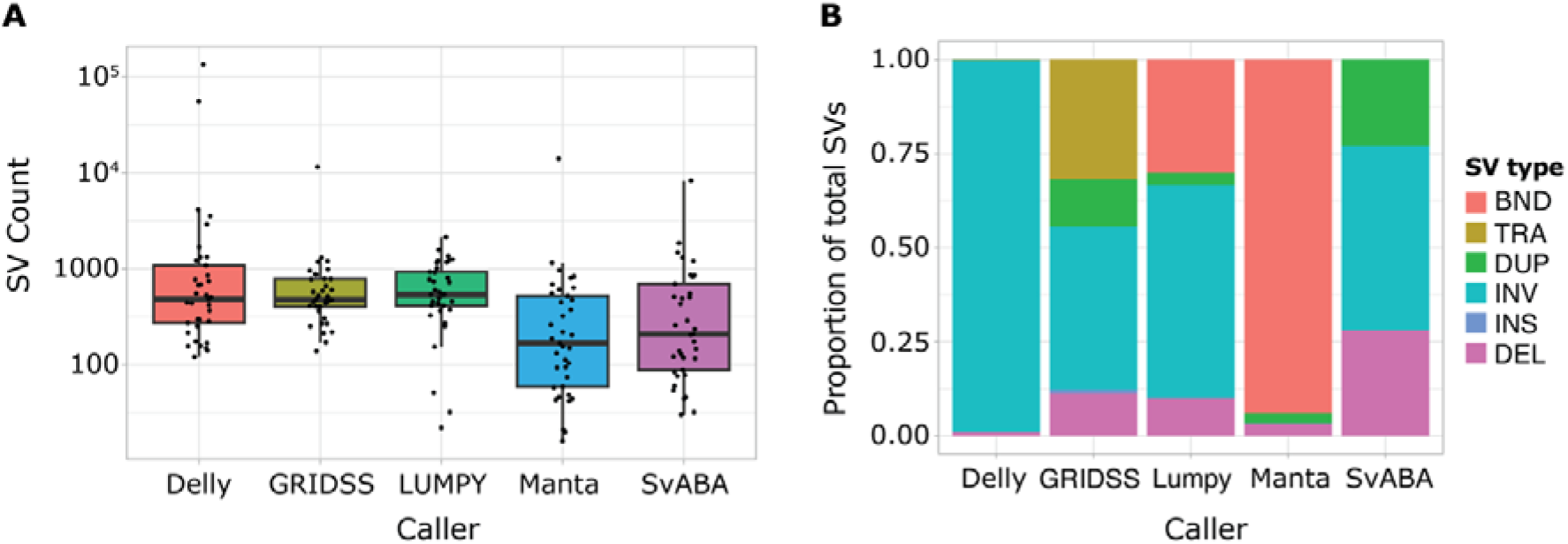
**(A)** The total number of SVs identified per sample by each caller in the validation cohort. **(B)** The proportions of each SV type identified by each caller. For comparison purposes, event types for GRIDSS and SvABA were annotated using publicly-available custom R scripts, as these callers identify breakends only. BND, breakend; TRA, translocation; DUP, duplication; INV, inversion; INS, insertion; DEL, deletion.

The five callers are known to identify different SV types (summarised in Table S1): Delly labels all event types; Manta and LUMPY call all events with the exception of inversions and insertions, respectively. GRIDSS and SvABA call breakends, without further labelling of variant types. To enable comparison between the callers, variants identified by SvABA and GRIDSS were assigned an event type using publicly-available specialised scripts (see Methods).

Comparing frequency of event types identified by these five callers, we found that the callers identify different proportions of each event type (Fig. 1B, Table S2). In this dataset, predominantly identified inversions (≥96% of events). Similarly, Manta primarily detected breakends (94% of events). GRIDSS and LUMPY showed more variation in the types of events identified, and called similar proportions of inversions (43% and 56%, respectively) and deletions (11% and 10%, respectively).

Structural variant calling is known to be resource-intensive^23^. We therefore assessed central processing unit (CPU) and memory usage to characterise the compute resource usage by each caller using the default settings. The five algorithms showed wide variation in both CPU and memory usage (Fig. S1). GRIDSS had the highest CPU usage, with a median of 39.3 CPU hours per sample (Fig. S1A). In contrast, Delly had the shortest median CPU time (4.1 hours). GRIDSS also showed the highest mean memory usage (median 12.5 GB; Fig. S1B); mean memory usage by SvABA was similar (median 10 GB). Of the five callers, Manta had the lowest memory usage; almost two orders of magnitude less than GRIDSS and SvABA (median 183 MB).

### Overlap with validation cohort calls

To benchmark the performance of individual callers, we compared SVs called by each tool with those from a validation set. This comprised variants from the 40 PCAWG patients profiled above, identified using a consensus approach of variants called by a least two of four tools (BRASS, Delly, SvABA and dRanger^20^). Across all samples, the validation set contained a total of 11,521 calls.

The overlap of calls made using Delly, GRIDSS, LUMPY, Manta and SvABA with the validation set was determined using a refined implementation of the PCAWG consensus calling approach. The highest proportion of validation set calls were identified by Manta (*n* = 10,123; 88%; Fig. 2). SvABA identified the next highest fraction (*n* = 9,287; 81%). Twenty-eight percent (*n* = 3,223) of the validation set calls were identified by all 5 callers. Fewer than 10% (*n* = 1,002) of PCAWG calls were identified by one caller only; SvABA was most likely to call variants not identified by another caller (510/1,002). A total of six hundred and forty-three (6%) of the validation set calls were not identified by any included caller.

**Figure 2.**
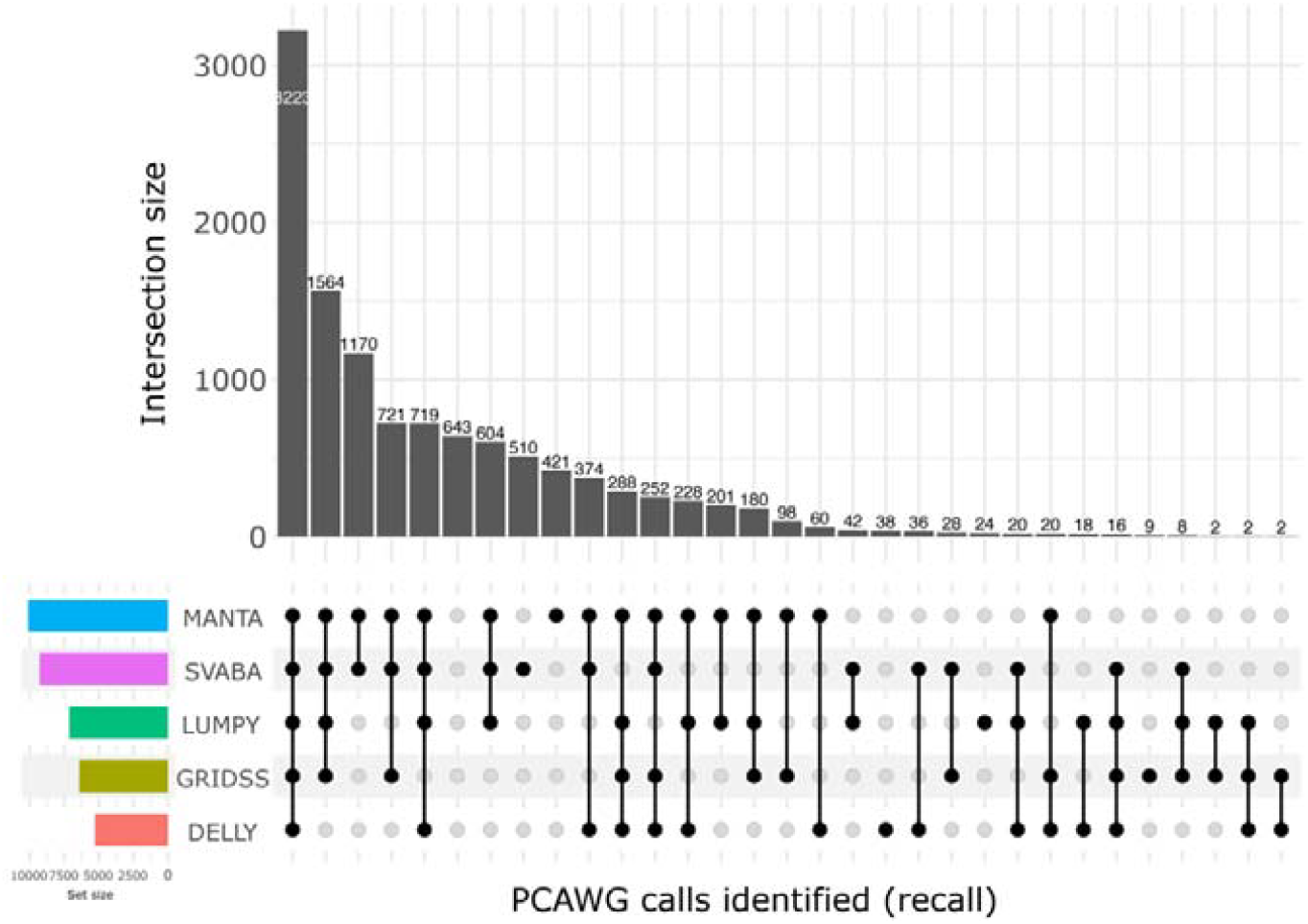
UpSet plot showing the number of validation set SV calls identified by each caller.

All callers except GRIDSS (30%) made high fractions of calls in addition to the validation set, although GRIDSS showed a lower recall in this series (Figure 2, Supplementary figure 2). Delly showed the highest fraction of extra calls, with 98% of its calls in excess of the validation set. LUMPY, Manta and SvABA made variable proportions of additional calls (73%, 63%, and 58% of total calls, respectively).

### A combination of four callers gives best overall performance

We next evaluated the performance of ensemble calling with a consensus of ≥2/*n* callers, using the validation set of 40 samples^20^. We evaluated two metrics: sensitivity (the fraction of calls identified) and proportion of calls made in excess of validation set. These values were calculated for all permutataions of caller combination; best performance was assessed using a balance between these two metrics.

Manta was consistently included in the best-performing combination in each SV category (Figure 3). Use of two-caller combinations (*i*.*e*. SVs must be identified by both callers to be included) showed relatively poor sensitivity compared to other approaches (Supplementary Figure 2. Of these, the combination of Manta and Delly gave the best performance, identifying 67% of validation calls while displaying 36% additional calls.

**Figure 3.**
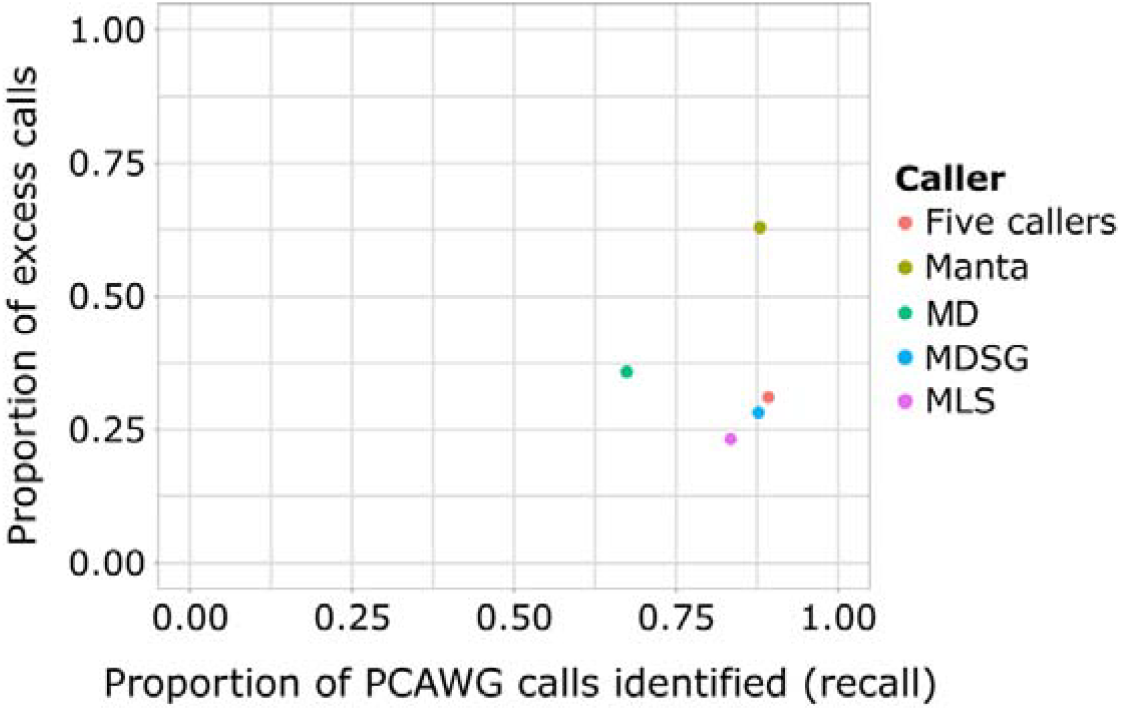
Scatterplot showing the best performing approach from each class of individual callers, and 2-, 3-, 4- and 5-caller combinations. Performance was determined using a balance between the highest proportion of validation set calls identified in combination with the lowest proportion of calls in excess of the validation set.

Among the three caller combinations, those including Manta and SvABA showed the highest recall (Supplementary figure 2, with Manta/SvABA/GRIDSS and Manta/LUMPY/SvABA featuring 80% and 83% recall, respectively). The majority of three caller combinations showed broadly similar proportions of calls in excess of the validation set (13-25%), with the exception of Manta/Delly/LUMPY (38%). Delly, GRIDSS, Manta and SvABA together showed the best performance of the four-caller combinations, identifying 88% of validation set calls with a proportion of 28% excess calls (Supplementary figure 2). Addition of LUMPY to this approach did not identify significantly more validation set calls (*p* = 0.11), but did increase computational time (median increase 8.2 CPU hours; *p* = 0.008) (Supplementary figure 3). The four-caller approach with Delly, GRIDSS, Manta and SvABA identified a similar fraction of validation calls to Manta, but more than halved the proportion of additional calls. This combination was therefore selected as best combination of sensitivity, minimised excess calls and CPU usage. A summary of the recall and excess call proportion for the best-performing approach in each category is in Table S3. The four-caller approach has been made available as a Nextflow pipeline, which we believe represents a valuable resource for the cancer genomics community.

## Discussion

Here, we report benchmarking of SV callers using patient-derived data. We ran five algorithms with individual documented precision to identify somatic variants, aiming for high sensitivity with minimal excess calls. In keeping with previous reports, Manta performed well as an individual caller^2^, and was included in the best-performing combinations in each class. A ≥2/4 consensus approach using Delly, GRIDSS, Manta and SvABA provided the best combination of sensitivity, minimised excess calls, and optimised CPU usage. This consensus approach, including panel of normal and somatic filtering, is available as a NextFlow pipeline (see Code availability).

The optimal combination of tools is likely to show some variation across datasets^9^. In this study, we determined the optimal number of callers to be four. In contrast, ConsensusSV achieves a similar validation rate using eight callers^14^. Increasing the number of callers has the potential to increase the rate of false positive calls. We observed no significant increase in recall when adding an additional caller (LUMPY) to the best-performing combination of four callers. Given the increased CPU time required and the potential to increase false positive calls, we propose implementation of the four-caller approach above.

There are limitations to this study. First, for comparison purposes, breakends called by SvABA and GRIDSS were annotated using custom scripts. The function used to annotate SvABA breakends identifies inversions, deletions and duplications/insertions only. The frequencies of breakends or translocations, and the relative frequencies of duplications and insertions are therefore unknown. Second, this study uses patient-derived data and estimates false positive calls using comparative metrics. It is possible that not all excess variants identified represent false positive calls, as some may be true SVs not present in the validation dataset. Although Delly and SvABA were included in both the approach described here and that used to generate the validation call set, Manta (which was not included in the PCAWG consensus), demonstrated the strongest performace in this analysis.

There is currently a lack of gold standard datasets for benchmarking real-world somatic structural variant calling. Here we used a set of consensus SVs generated by a different study with a different combination of callers, with additional evaluation of specificity and proportion of additional calls using updated tools. Although the best-performing individual caller (Manta) and four-caller consensus showed similar precision, use of the consensus approach more than halved the proportion of excess calls. Thus, in this dataset, use of optimised consensus calling improved accuracy while maintaining sensitivity. However, it is possible in other settings that the use of a consensus approach may increase accuracy at the expense of sensitivity^5^.

Our optimised consensus SV calling approach and subsequent analysis provide an important contribution to the cancer genomics community. A practical implementation of this method has been made available as a NextFlow pipeline. Although individual callers such as Manta show strong standalone performance, use of a consensus strategy minimises excess calls, reinforcing the value of consensus calling for accurate SV identification. Similar consensus approaches, such as FusorSV^17^ and MetaSV^16^ lack more recent tools or an assembly-based caller, which has been recommended in benchmarking studies using cell lines^9^. These tools, along with ConsensusSV^14^, have included at least one ‘imprecise’ caller (*i*.*e*. precision <30%), which worsens the precision of consensus call sets^2^. Here, we determine the optimal combination, set expectations for the expected SV burden and lay a foundation for further development and refinement of SV calling tools.

## Methods

Binary Alignment/Map (BAM) files for a validation cohort (*n* = 40) patient-matched tumour and germline samples from the TCGA portion of the Pan-Cancer analysis of Whole Genomes Consortium^20^ with were realigned from hg19 to hg38 using DRAGEN^24^. In all cases, human leucocyte antigen sequences, alternate and decoy contigs were excluded.

SVs were identified from the realigned BAM files using five specialised callers: Manta^18^ (v1.5), Delly^21^ (v0.8.5), LUMPY^25^ (implemented as Smoove v0.2.6), SvABA^22^ (v1.1.0) and GRIDSS^19^ (using the gridss-purple-linx suite, v1.3.2). Callers were implemented with default settings. The minimum SV size detected varies by caller: by default, Manta assembles SVs 8 base pairs (bp) or larger, whereas GRIDSS defaults to 10 bp. SvABA and LUMPY have a lower size detection limit of 50 bp and 100 bp, respectively. The minimum SV size called by Delly varies according to insert size distribution. For an insert size of 200-300 bp with a standard distribution of 20-30 bp, Delly has a reliable lower detection limit of 300 bp.

The resulting Variant Call Format (VCF) files were filtered using BCFtools^26^ (v1.12) to remove variants present in the matched germline sample, with the exception of Delly. For this tool, somatic variants were identified using *delly filter*, for which a minimum alternate allele fraction support of was used (as recommended by the developers).

BCFtools was used to filter germline VCF files to retain germline-only variants, with *bcftools filter* -*i ‘GT[0]=“alt*”‘; imprecise variants were removed using *grep*. A panel of normals was then generated on a caller-specific basis with these variants, using a custom script. This annotates each variant present in the germline VCF files with the frequency with which it occurs across all germline samples. For GRIDSS, somatic filtering was performed using GRIDSS Post Somatic Software (GRIPSS, version 1.9). Using a panel of normal v- ariants generated for this cohort, somatic SVs present in ≥4 samples were flagged as panel of normal variants and removed.

Following panel of normal and somatic filtering, a modified version of a previously-described approach^20^ was used to generate consensus somatic SVs identified by individual callers and ≥2/*n* of 2, 3, 4 and 5 callers. Somatic SVs generated using a consensus approach comprising a different combination of callers^20^ were available for the validation cohort. These variants were lifted over from hg19 to hg38 using UCSC LiftOver^27^ (version 1.0). A modified version of the approach above was then used to identify somatic SV calls present in both the test and validation call sets for each caller individually, and in combinations of 2, 3, 4 and 5.

SvABA and GRIDSS are breakend callers. To enable direct comparison of identified event types, event types for SvABA and GRIDSS were assigned using specialised scripts. Variants identified by SvABA were annotated using the annotate_sv_type function from the variantCall^28^ R package. The simple_event_type.R script provided by the developers^29^ was used to assign variant types to calls made by GRIDSS.

All descriptive statistics were calculated using R (version 4.0.2). Sensitivity was calculated as the proportion of calls in the validation set identified by each caller or combination of callers. The proportion of excess calls identified by each approach was calculated as the number of calls made in addition to the validation call set, as a proportion of the total calls made by each approach.

## Acknowlegements

S.W. was supported by a Jean Shanks/Pathological Society Clinical Lecturer (JSPS CLG 2019 01) and a Pathological Society Trainees’ Small Grant (TSGS 0421 1297). S.W., N.M., T.L., J.D. and P.V.L. were supported by the Francis Crick Institute, which receives its core funding from Cancer Research UK (CC2008), the UK Medical Research Council (CC2008) and the Wellcome Trust (CC2008). J.D. was supported by a postdoctoral fellowship from the Research Foundation – Flanders (project no. 12J6916N) and acknowledges current grant support from VIB (Vlaams Instituut voor Biotechnologie). N.P. holds a Cancer Research UK Career Establishment Award (ref: RCCCEA-Nov23/100003). Support was provided to N.P. and A.M.F. by the National Institute for Health Research, the University College London Hospitals Biomedical Research Centre, and the Cancer Research UK University College London Experimental Cancer Medicine Centre. P.V.L. is a CPRIT Scholar in Cancer Research and acknowledges CPRIT grant support (RR210006).

## Code availability

A NextFlow implementation of the described consensus approach is available in the waise_consensus repository (https://github.com/sarawaise/waise_consensus).

## Ethics declarations

The authors declare no competing interests.

