## Supplementary Information for "An optimised computational approach for the identification of somatic structural variants in cancer"

**SUPPLEMENTARY INFORMATION	1**

**Supplementary figures	2**

**Supplementary tables	4**

**Supplementary Figures**

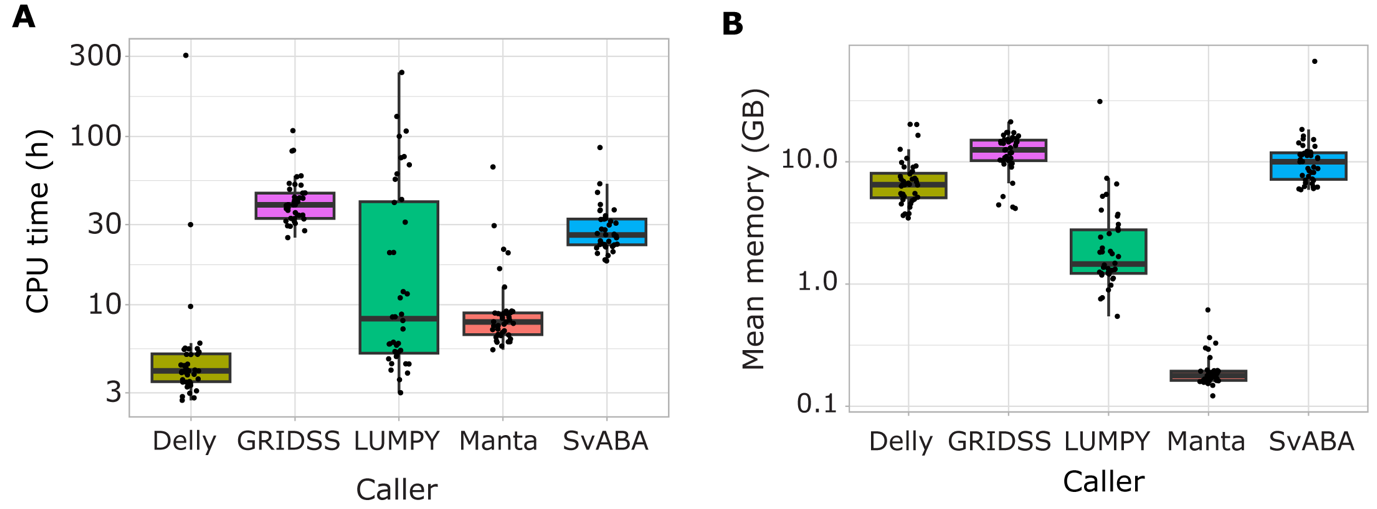
**Supplementary Figure 1.** (**A**) CPU and (**B**) mean memory usage *per* validation set sample for each caller. Y-axis log scale.

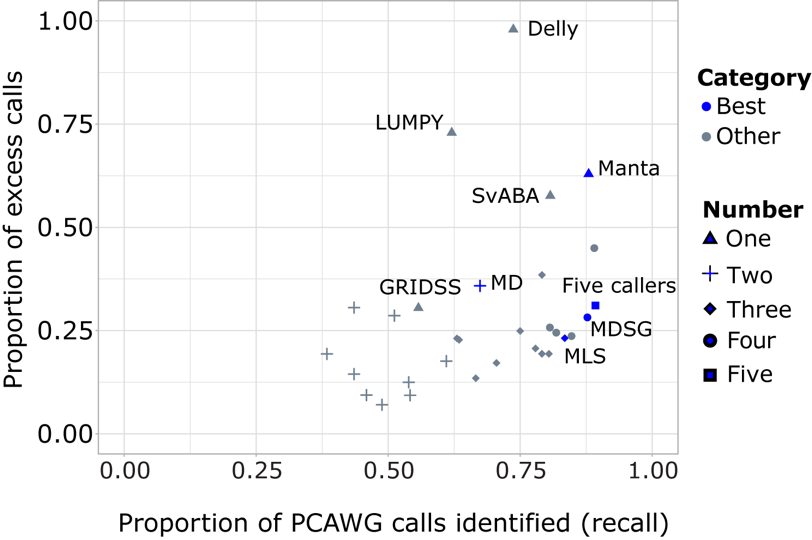

**Supplementary Figure 2.** Scatterplot showing the best performing approach from each class of individual callers, and 2-, 3-, 4- and 5-caller combinations. Performance was determined by the highest proportion of validation set calls identified in combination with the lowest proportion of calls in excess of the validation set. MD, Manta and Delly; MLS, Manta, LUMPY and SvABA; MDSG, Manta, Delly, SvABA and GRIDSS.

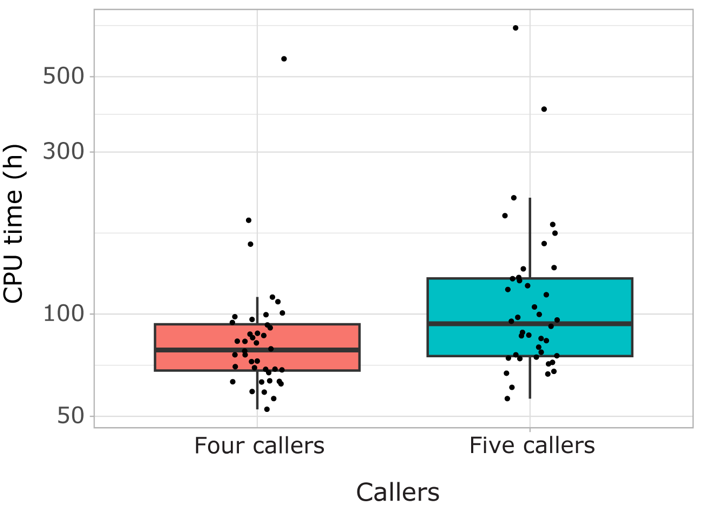

**Supplementary figure 3.** Boxplot showing the central processing unit (CPU) time *per* sample for a combination of Delly, GRIDSS, SvABA and Manta, and this combination with the addition of Manta. Y-axis log scale.

**Supplementary Tables**

|  | **BND** | **DEL** | **DUP** | **INS** | **INV** |
| --- | --- | --- | --- | --- | --- |
| Manta | x | x | x | x |  |
| Delly | x | x | x | x | x |
| LUMPY | x | x | x |  | x |
| SvABA | x |  |  |  |  |
| GRIDSS | x |  |  |  |  |

**Table S1.** Summary of the structural variant types identified by each caller.

|  | **BND** | **DEL** | **DUP** | **INS** | **INV** | **TRA** |
| --- | --- | --- | --- | --- | --- | --- |
| Manta | 94.0 | 3.2 | 3.7 | 0.1 | 0 | 0 |
| Delly | 0.1 | 1.0 | 0.2 | 0 | 98.7 | 0 |
| LUMPY | 30.2 | 10.0 | 3.3 | 0 | 56.5 | 0 |
| SvABA | 0 | 2.2 | 1.8 | 0 | 96.0 | 0 |
| GRIDSS | 0 | 11.1 | 3.3 | 1.2 | 43.4 | 31.8 |

**Table S2.** Proportions of event types identified by each caller. For comparison purposes, event types for GRIDSS and SvABA were annotated using custom R scripts, as these callers identify breakends only.

| **Category** | **Caller or combination of callers** | **Sensitivity (%)** | **Excess calls (%)** |
| --- | --- | --- | --- |
| One caller | Delly | 74 | 98 |
|  | GRIDSS | 56 | 30 |
|  | LUMPY | 62 | 73 |
|  | **Manta** | **88** | **63** |
|  | SvABA | 81 | 58 |
| Two callers | Delly/GRIDSS | 44 | 14 |
|  | Delly/LUMPY | 51 | 29 |
|  | Delly/SvABA | 44 | 31 |
|  | LUMPY/GRIDSS | 46 | 9 |
|  | LUMPY/SvABA | 54 | 12 |
|  | **Manta/Delly** | **67** | **35** |
|  | Manta/GRIDSS | 54 | 9 |
|  | Manta/LUMPY | 61 | 18 |
|  | Manta/SvABA | 39 | 19 |
|  | SvABA/GRIDSS | 49 | 7 |
| Three callers | Delly/LUMPY/GRIDSS | 63 | 23 |
|  | Delly/SvABA/GRIDSS | 53 | 23 |
|  | Delly/LUMPY/SvABA | 75 | 25 |
|  | LUMPY/SvABA/GRIDSS | 67 | 13 |
|  | Manta/Delly/LUMPY | 79 | 38 |
|  | Manta/Delly/GRIDSS | 78 | 21 |
|  | Manta/Delly/SvABA | 79 | 27 |
|  | **Manta/LUMPY/SvABA** | **83** | **23** |
|  | Manta/SvABA/GRIDSS | 80 | 19 |
|  | Manta/LUMPY/GRIDSS | 71 | 17 |
| Four callers | Delly/LUMPY/SvABA/GRIDSS | 81 | 26 |
|  | Manta/Delly/LUMPY/GRIDSS | 82 | 24 |
|  | Manta/Delly/LUMPY/SvABA | 89 | 45 |
|  | **Manta/Delly/SvABA/GRIDSS** | **88** | **28** |
|  | Manta/LUMPY/SvABA/GRIDSS | 85 | 24 |
| All callers | **N/A** | **89** | **31** |

**Table S3.** The sensitivity and the calls in excess of the validation set as a percentage of total calls identified by the best-performing approach in each category tested. The best-performing caller, or combination of callers, in each category is highlighted in bold.
